# Transcriptional dynamics following freezing stress reveal selection for mechanisms of freeze tolerance at the poleward range margin in the cold water intertidal barnacle *Semibalanus balanoides*

**DOI:** 10.1101/449330

**Authors:** Katie E. Marshall, Eddy J. Dowle, Alexandra Petrunina, Gregory Kolbasov, Benny K. K. Chan

**Author notes:** Corresponding author: Benny K.K. Chan.

## Abstract

The ability to survive freezing has repeatedly evolved across multiple phyla. This suggests that the mechanisms of freeze tolerance must be readily evolvable from basal physiological traits. While several biochemical correlates to freeze tolerance have been described, the mechanism that confers freeze tolerance is still not well understood. To understand both the basic biochemical mechanisms of freeze tolerance as well as their role in local adaptation at the poleward range edge, we conducted a transcriptomic study on two populations (one from the poleward range margin in the White Sea, Russia, and one from the central coast of British Columbia, Canada) of the cold water acorn barnacle *Semibalanus balanoides* on a time series following a freezing event. We found that the British Columbia population (at the equatorward range margin) was significantly less freeze tolerant than the White Sea population (at the poleward range margin). After assembling and annotating a *de novo* transcriptome for *S. balanoides*, we found that the patterns of differential transcript expression following freezing were almost entirely non-overlapping between the two populations, with the White Sea population expressing a series of heat shock proteins in response to freezing stress as well as several aquaporins, while the British Columbia population expressed a series of proteases instead, indicating severe protein damage. We found strong evidence of purifying selection on the significantly upregulated transcripts in the White Sea population, suggesting local adaptation to freezing threat. Taken together, this shows the importance of freeze tolerance to population survival at the poleward range margin, and highlights the central roles of aquaporins and heat shock proteins to the trait of freeze tolerance across taxa.

## Introduction

The occurrence of species range edges is one of the central problems in biogeography (Sexton, Mcintyre, Angert, & Rice, 2009). Broadly speaking, it is believed that equatorward range edges are due to biotic interactions, while poleward range edges are due to abiotic effects (Louthan, Doak, & Angert, 2015). Given that minimum temperatures have the steepest and most consistent latitudinal gradient (Sunday, Bates, & Dulvy, 2012), this suggests that low temperature tolerance is central for understanding the presence of poleward range edges. Therefore, understanding the mechanisms and adaptive potential of low temperature tolerance is central for delineating the role of local adaptation in the maintenance of poleward range edges.

Low temperature tolerance includes responses to two overlapping, but distinct, physiological stressors: the direct effects of low temperature on reaction rates, and managing the potential and spread of internal ice. In ectothermic organisms, biochemical reaction rates decrease due to Arrhenius effects as well as by decreased membrane fluidity at low temperatures, which causes a general decrease in rates of physiological functioning (reviewed in R. E. Lee, 2010). At body temperatures below 0 °C, organisms also risk internal ice formation. Surviving these two stressors, while linked by temperature, appear to involve different physiological mechanisms. Some organisms like the eastern spruce budworm (*Choristoneura fumiferana* Clemens) are able to suppress their freezing point well below −30 °C by the production of low molecular weight cryoprotectants and antifreeze proteins, and can survive low temperatures as long as they do not freeze (Han & Bauce, 1995; Katie E. Marshall & Sinclair, 2015; Qin, Doucet, Tyshenko, & Walker, 2007). Others like the woolly bear caterpillar (*Pyrrharctia isabella* Smith) induce and survive freezing at relatively high temperatures (c.a. −10 °C), but are unable to survive temperatures below −16 °C (Boardman, Terblanche, & Sinclair, 2011; Layne & Kuharsky, 2000). Being able to survive freezing is therefore a physiological trait that is likely mechanistically distinct from being able to survive low temperature (reviewed in Toxopeus & Sinclair, 2018).

The ability to survive internal ice formation is a paraphyletic trait broadly distributed across the animal kingdom, with freeze-tolerant species from multiple phyla including arthropods, molluscs, and vertebrates (Sinclair, Addo-Bediako, & Chown, 2003; Storey & Storey, 1996; Toxopeus & Sinclair, 2018). Freeze tolerance appears to have evolved independently on at least 20 occasions, and this is almost certainly a large underestimate (Sinclair et al., 2003). The repeated emergence of this trait suggests that the underlying mechanisms must be readily evolvable and that freeze tolerance is possible in a variety of underlying physiological conditions. Despite the ubiquity of the trait, and over 200 years of research, the physiological mechanisms underlying animal freeze tolerance remain poorly understood (Lee, 2010; Sømme, 2000; Teets & Denlinger, 2014; Toxopeus & Sinclair, 2018).

Surviving internal ice formation is a unique challenge for animals. They need to be able to manage potential mechanical damage from ice crystal formation, cellular dehydration as water leaves the cell to join the growing extracellular ice lattice, and sustained periods of anoxia since oxygen cannot perfuse easily to frozen tissue (Lee, 2010). Many biochemical correlates to survival of these challenges have been described; most commonly this involves the mass biosynthesis of low molecular weight polyols and sugars that act as osmoprotectants and cryoprotectants that can also decrease the melting (and therefore freezing) point of cells for avoiding intracellular ice formation (Koštál, Korbelová, Poupardin, Moos, & Šimek, 2016; Toxopeus & Sinclair, 2018). However there is no correlation between quantity of low molecular weight cryoprotectants and degree of freeze tolerance—for instance, woolly bear caterpillars *P. isabella* accumulate circa 2.5 M glycerol as a cryoprotectant but have a lower lethal temperatures of only circa −16 °C (Boardman et al., 2011; K. E. Marshall & Sinclair, 2011), while the freeze tolerant goldenrod gall fly (*Eurosta solidaginis* Fitch) has glycerol content of circa 0.3 M but a lower lethal temperature below −50 °C (Lee, Dommell, Joplin, & Denlinger, 1995). Freeze tolerant intertidal barnacles and mussels appear to not accumulate polyols or sugars at all (Johnston & Clarke, 1990). It therefore appears that while these biochemical correlates might be associated with enhanced survival of freezing, they do not explain the physiological mechanisms of the ability to tolerate internal ice.

More recent work has focused on the role of proteins in freeze tolerance. Antifreeze proteins may protect against intracellular freezing, which is almost always lethal even in very freeze tolerant animals (Davies, 2014; but see review in Sinclair & Renault, 2010). In the extracellular spaces, they may inhibit recrystallization, keeping ice crystals small enough to avoid mechanical damage. However they have been described from only a few freeze tolerant species, and their role in freeze tolerance remains controversial (Duman, Bennett, Sformo, Hochstrasser, & Barnes, 2004; Toxopeus & Sinclair, 2018). Aquaporins facilitate the movement of water out of the cell during freezing and glycerol transporters to facilitate the movement of sugars into the cell have been reasonably well-explored in plant freeze tolerance (Peng, Arora, Li, Wang, & Fessehaie, 2008; Philip, Yi, Elnitsky, & Lee, 2008; Storey & Storey, 2013; Yi et al., 2011). In animals, aquaporins are implicated in the freeze tolerance in *E. solidaginis* and the wood frog (*Lithobates sylvatica* LeConte), but how taxonomically broadly important they are for freeze tolerance is not currently well understood (Philip, Kiss, & Lee, 2011; Philip et al., 2008; Storey & Storey, 2013; Yi et al., 2011). Finally, heat shock proteins are constitutively upregulated in overwintering insects and have been implicated in plant freeze tolerance, but are not generally thought to be involved in responses to acute freezing stress in animals (King & MacRae, 2015; Rinehart et al., 2006). However, the Antarctic midge (*Belgica antarctica* Jacobs) upregulates Hsp70 in response to repeated freezing episodes, and freeze tolerant drosophilid fly (*Chymomyza costata* Zetterstedt) upregulate small heat shock proteins following freezing suggest, suggesting that heat shock proteins may be more important to freeze tolerance than previously thought (Teets, Kawarasaki, Lee, & Denlinger, 2011).

While the biochemical correlates of freeze tolerance have been explored in broad array of terrestrial organisms, very little is known about the biochemical mechanisms of freeze tolerance in intertidal organisms (Dennis, Loomis, & Hellberg, 2014; Murphy, 1983). In temperate and polar regions, intertidal organisms including barnacles, mussels, and snails are almost without exception freeze tolerant (Murphy, 1983; Waller, Worland, Convey, & Barnes, 2006; but see Sinclair, Marshall, Singh, & Chown, 2004). This is necessary because of the unique nature of the intertidal—at low tide in the winter, the wetness of the habitat and the year-round presence of food in the digestive tract makes freezing a constant threat when animals are emersed. Some efforts have been made to determine the underlying biochemical mechanisms, but they remain elusive (Ansart & Vernon, 2003; Dennis et al., 2014; Murphy, 1983; Storey & Storey, 2013). It has been hypothesized that anaerobic byproducts such as strombine and taurine may be important in freeze tolerance of *Mytilus sp.* mussels (Loomis, Carpenter, & Crowe, 1988), but much remains unknown about how intertidal species survive freezing.

Here we present the first (to our knowledge) transcriptomic study following freezing stress in an intertidal animal. We explore potential mechanisms of freeze tolerance in the acorn barnacle (*Semibalanus balanoides* Linnaeus) from two populations that differ widely in the intensity of freezing stress, one from the central coast of British Columbia, Canada which experience relatively mild winters, and one from the White Sea in Russia which experiences significantly harsher winters. We hypothesize that barnacles from British Columbia will be less freeze tolerant (will have lower survival following freezing) than barnacles from the White Sea. If this is true, we will infer that transcriptional responses to freezing in the White Sea population likely enhance freeze tolerance. In addition, if transcripts upregulated in this population are important for survival in poleward range edge habitats, we predict that we will see evidence of selective sweeps on transcripts identities that are associated with biochemical mechanisms of freeze tolerance and are upregulated following freezing stress. We find strong interpopulation differences in response to freezing stress, and that this includes wholescale transcription of heat shock proteins following freezing stress, as well as the transcription of aquaporins (which show significant evidence of selective sweeps) and potentially new classes of antifreeze proteins. This suggests that these protein-based mechanisms of freeze tolerance may be broadly universal across freeze tolerant phyla, and that they may have selective advantages in intertidal environments.

## Methods

### Study system and collection

The barnacle *S. balanoides* is a common species in rocky intertidal zones through the Arctic Circle, including northwestern Europe and both coasts of North America. In western North America, it is found as far south as the central coast of British Columbia (Flight & Rand, 2012; Schmidt, Bertness, & Rand, 2000; Wethey, 1983). Because of its polar distribution as well as its intertidal habitat, *S. balanoides* regularly experiences freezing events throughout the winter (Cook & Gabott, 1970; Cook & Lewis, 1971; Dennis J. Crisp & Ritz, 1967). Late-stage larvae (cyprids) of *S. balanoides* have even been found embedded in ice during winter, and successfully settle and metamorphose after spring thaw (Pineda, DiBacco, & Starczak, 2005). Barnacles do not appear to use polyol cryoprotectants for their freeze tolerance, but little is known about the mechanisms of their freeze tolerance (Cook & Gabott, 1970; D.J. Crisp, Davenport, & Gabbott, 1977).

We collected *S. balanoides* from mid intertidal habitats by gathering small stones covered with large groups of barnacles at the White Sea Biological Station (Moscow State University, Russia; 66.5551° N, 34.1976° E) as well as from Calvert Island, British Columbia, Canada (51.5559° N, 128.0365° W) during the spring of 2015. The climate on Calvert Island is moderated by the Alaska Current which brings warm water from the south, and thus winter temperatures rarely drop below −10 °C, while in the White Sea overwinter temperatures regularly drop below −20 °C (Figure S1-S2), thus we expected barnacles from the White Sea be more tolerant of freezing. We have installed robobarnacles (Chan, Lima, Williams, Seabra, & Wang, 2016) in the White Sea to record the estimated body temperature of similarly-shaped *Tetraclita spp*. barnacles during the winter months (Figure S1). Barnacles for transcriptomic experiments were collected in June in both habitats, wrapped in wet newspaper, then were placed in a common seawater aquarium maintained at 10 °C and aerated by an airstone for two weeks before experimentation.

### Experimental design and freezing exposures

After acclimation, we measured barnacle test diameter with digital calipers along the longest edge to the nearest mm. To simulate an intertidal freezing event, we cooled barnacles by placing the rocks they had settled on into a Styrofoam box, then placed the box in a freezer set to −20 °C. We continuously monitored barnacle temperature by placing a 36 AWG gauge Type T (copper-constantan) thermocouple through the plates into the barnacle body. The thermocouple was attached to a Picotech TC-08 interface (Pico Technology, Cambridge UK) which relayed temperature data to an attached computer every 0.5 s in real time. We maintained barnacle temperature to within 0.5 °C of the desired temperature by opening and closing the freezer door. Barnacle supercooling points (SCPs) were recorded as the lowest temperature attained before sudden release of heat due to the latent heat of crystallization (Sinclair, Coello Alvarado, & Ferguson, 2015). With this procedure, we achieved cooling rates of circa 1.2 °C/min, which is within the range found in their natural habitat during a low tide (Figure S1). To measure differences in freeze tolerance, we exposed barnacles from both populations to −10 or −6 °C for 4 h, then warmed them by placing them immediately into seawater at 10 °C to mimic inundation during high tide conditions. Survival was monitored for two days following and scored based on the ability to close the top plates when removed from seawater, then compared using generalized linear models with a binomial distribution in R (Team, 2016), while SCPs were compared using ANOVA.

To measure transcriptional changes following freezing, we exposed 48 barnacles from each population to −6 °C for 4 h using the above procedure. We sampled groups from each population at four different time points: before the freezing event took place (“control”), after the freezing event but before thawing in seawater (“before thawing”), 4 h after thawing (“4h recovery”) and 28 h after thawing (“28h recovery”; experimental design presented in Figure 1). For each population × timepoint, we rapidly dissected out the soft tissue of 12 barnacles from each population and flash-froze tissue in groups of three individuals in liquid nitrogen for four total biological replicates/population/timepoint.

**Figure 1.**
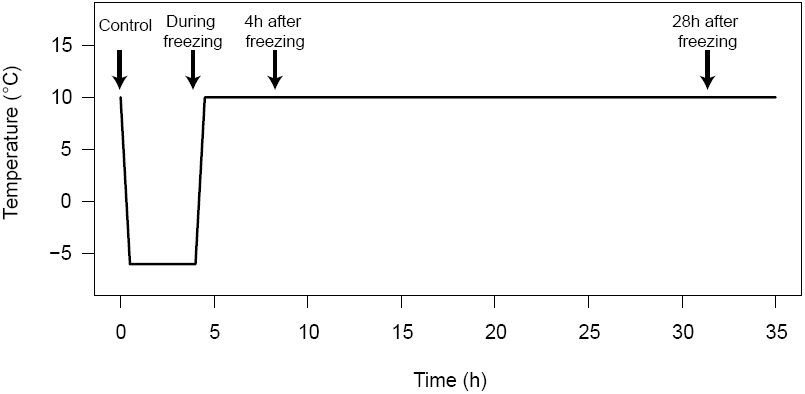
Outline of experimental design, showing timepoints that barnacles from each population were sampled.

### Transcriptomics

Total RNA was extracted from 32 pools of barnacles using a TRIzol protocol. Sequencing libraries were prepared and sequenced by the Academia Sinica Bioinformatics Center. Libraries were prepared using Illumina’s stranded mRNA TruSeq RNA sample prep v2 kit as per manufacturer’s instructions. Cleaned libraries were sequenced on 2 lanes of 181 bp paired-end Illumina HiSeq 2000 at the Academia Sinica Sequencing Center.

Our general approach for data analysis was informed by best practices for RNA-seq (Conesa et al., 2016). The preliminary data quality checks and analysis were conducted in Galaxy (Afgan et al., 2016). We clipped Illumina adapters from reads and removed low quality and short read sequences using Trimmomatic’s default settings (version 0.36.3; Bolger, Lohse, & Usadel, 2014). We then took the trimmed reads and assembled a transcriptome using Trinity (default settings for paired end reads, version 2.4.0, Grabherr et al., 2011), then clustered redundant sequences using CD-HIT-EST with a sequence identity cut-off of 0.9 (Fu, Niu, Zhu, Wu, & Li, 2012). We then aligned these sequences to the NCBI NR database using Diamond (Buchfink, Xie, & Huson, 2014), then filtered out all reads that did not map to metazoans (viruses, algae, bacteria, etc.) using MEGAN6 (Huson et al., 2016) to produce a *de novo* transcriptome for *S. balanoides*. To annotate the transcriptome, we used Blast2GO (default settings, version 5.1.13, Gotz et al., 2008) to blastx the transcripts to the NCBI arthropod nr database, then mapped to Interpro IDs and GO terms to produce a final, annotated transcriptome.

To quantify gene expression following a freezing event, we then mapped our trimmed reads to this transcriptome using HISAT2 (D. Kim, Langmead, & Salzberg, 2015), then assembled and quantified transcripts using StringTie (Pertea et al., 2015), gffCompare (Trapnell et al., 2010), and FeatureCounts (Liao, Smyth, & Shi, 2014). Following quantification, we tested for differences in transcript abundance among time points and populations using DESeq2 (Love, Huber, & Anders, 2014) to identify differentially expressed transcripts (adjusted p-value < 0.05) within each population as a function of timepoint, while controlling for multiple comparisons using the Benjamini-Hochberg multiple comparisons procedure.

We then used Blast2GO to test for functional enrichment of the significantly differentially regulated transcripts within each population. We simplified the significantly enriched GO list for plotting using REVIGO to eliminate semantically redundant terms, using the “tiny” setting (Supek, Bošnjak, Škunca, & Šmuc, 2011). We also annotated each sequence with the nearest KEGG orthology term using the KEGG Automatic Annotation Server (Moriya, Itoh, Okuda, Yoshizawa, & Kanehisa, 2007), then tested for KEGG pathway enrichment using the Gage and Pathview (Luo & Brouwer, 2013) packages in R.

### Population differentiation

To quantify genetic differences between the two geographic sites, single nucleotide polymorphisms (SNPs) were called using ANGSD 0.916 (Korneliussen, Albrechtsen, & Nielsen, 2014). The ‘before thawing’ samples were removed from the analysis as their transcript expression profiles and therefore individual transcripts read coverage levels, differed markedly from the other samples. Mapped reads were first filtered to remove PCR duplicates and secondary mapping. Within ANGSD missing values were assigned for base calls of a phred score <30. Loci were limited to those that had a depth of at least 10 reads per sample and an across all samples Minor Allele Frequency (MAF) estimate of >0.05. As each sample is comprised of three individuals (pooled) genotypes could not be called, instead sample MAF were estimated using the allele counts method in ANGSD. Sample MAF estimates were used to conduct a Principle Components Analysis (PCA) within R (prcomp function; PCA presented in Figure S3).

We tested whether there was evidence for selection on the differentially expressed transcripts at either of the two geographic sites. As with the MAF estimates, the ‘before thawing’ samples, PCR duplicates, and secondary mapping were removed. Samples were grouped into two groups by geographic site, with each of the two groups were comprised of 36 individuals total (12 samples of three individuals each). A pileup file was generated and Tajima’s D (Tajima, 1989) estimates were generated for both locations using PoPoolation (Robert Kofler et al., 2011). Within PoPoolation missing values were assigned for base calls of a phred score <20 and a corrected Tajima’s D was calculated for both geographic sites. Tajima’s D estimates were compared between the two sites within R (via a T-test). To further examine genetic differentiation between the two geographic sites. PoPoolation2 (R. Kofler, Pandey, & Schlotterer, 2011) was also used to estimate Fisher Exact Tests (per site) and F_ST_ values (per site and sliding window).

## Results

### Assessing freeze tolerance

Supercooling point (mean = −3.2 ± 0.51 °C) did not differ between barnacle populations, nor did SCP differ with basal diameter of test (p > 0.4 for both terms, n = 8 – 15 per population). Barnacles from the White Sea had significantly higher survival at both exposure temperatures (81 – 95% vs. 27 – 78%; Figure 2; deviance explained = 48.4, df = 1,18, p < 0.001), and exposure to −10 °C significantly decreased survival in both populations relative to exposure at −6 °C (deviance explained = 74.8, df = 1, 17, p < 0.001; n = 92 −152 for each combination).

**Figure 2.**
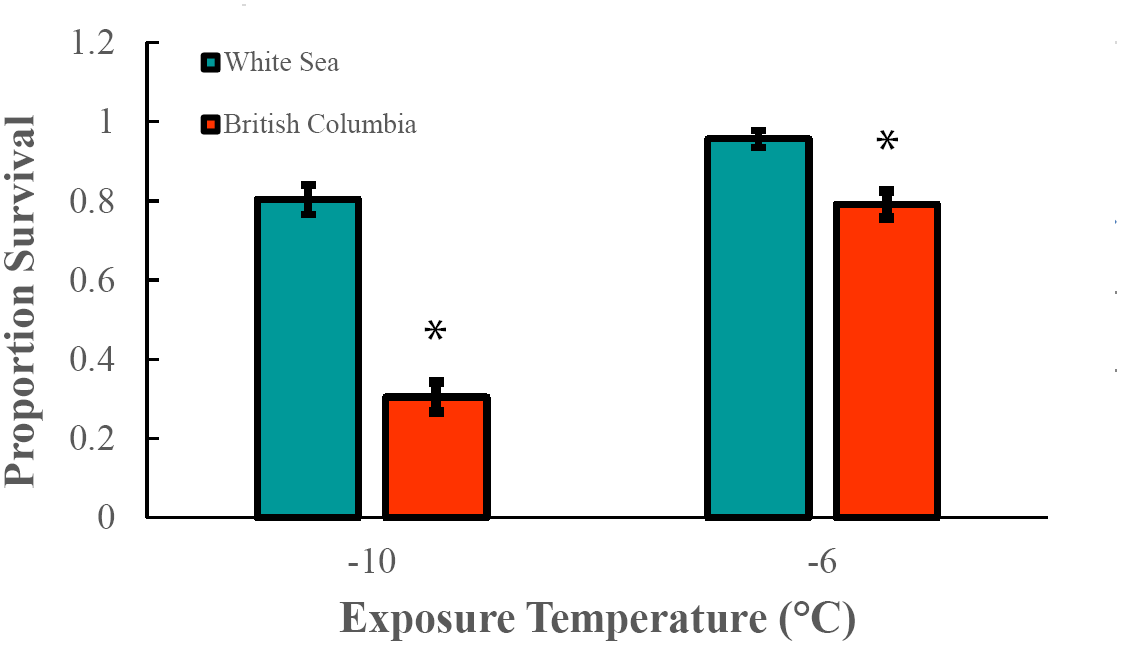
Survival of barnacles from the White Sea and British Columbia populations following a 4 h exposure to either −10 or −6 °C. Asterisk indicates a significant difference in survival between populations in a generalized linear model with binomial error, n = 92 – 151 per population/temperature.

### Sequencing and assembly

A total of 776.69 million 181 bp reads were produced from two lanes of Illumina Hiseq 2000, with an average of 24.75 million/library for the White Sea population and 23.79 million/library for the British Columbia population. Of these, an average of 14.2 million reads/library in the White Sea population and 13.1 million reads/library in the British Columbia population passed quality control by Trimmomatic. These reads were then used to assemble a *de novo* transcriptome using Trinity, which was then reduced to 61,881 sequences using CD-HIT-EST. Of these, a total of 61,779 were matched to taxonomic hits by Diamond. We then filtered nonmetazoan sequences (RNA viruses, algae, and other microorganisms) out using MEGAN, and were left with a total of 46,535 sequences that were an average of 1277 base pairs long (N_50_ = 1084 bp). An average of 4.84 million reads/library in the White Sea population and 3.63 million reads/library in the British Columbia population were mapped to the final transcriptome.

### Pattern of transcript expression

We found the patterns of significantly differentially expressed transcripts after a freezing event differed strongly between the two populations (Figure 3, Figure 4). In the more freeze tolerant population (White Sea), no transcripts were differentially expressed during freezing, while after four hours of recovery, 191 transcripts were differentially expressed (almost evenly split between up and downregulated). By 28 hours after freezing, this had decreased to 185 transcripts, with slightly more upregulated than down. This pattern was dramatically different in the less freeze tolerant population (British Columbia), which had a total of 11,050 transcripts differentially regulated during the freezing event, the majority of which (9432) were significantly upregulated. By four hours of recovery, this number had decreased to 51 (again dominated by 46 upregulated transcripts), and by 28 hours of recovery this had increased to 1739 transcripts (again dominated by 977 significantly upregulated transcripts).

**Figure 3.**
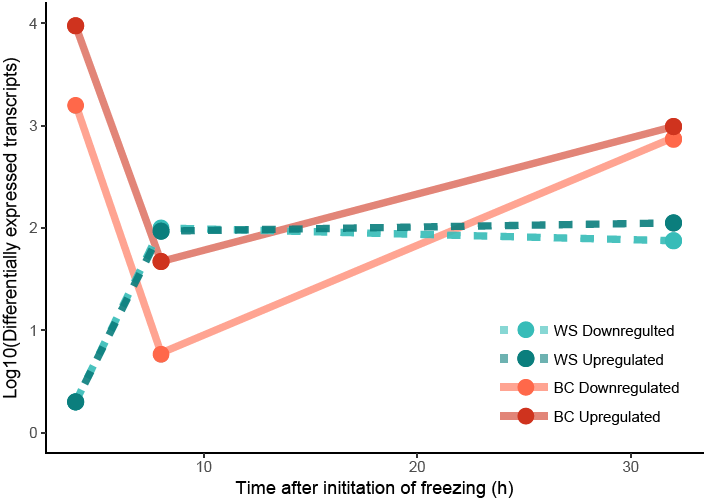
Timecourse of the number of differentially expressed transcripts in barnacles from the White Sea (WS) and British Columbia (BC) with expression calculated relative to control (0 h) for each population using DESeq2.

**Figure 4.**
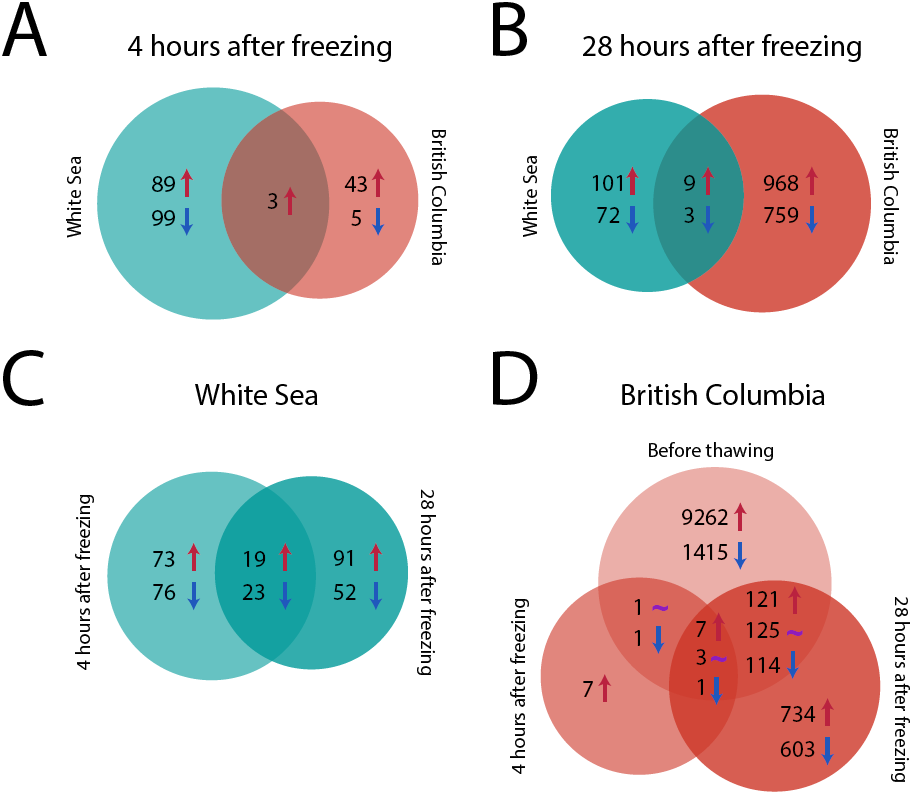
Venn diagrams illustrating the number of differentially expressed (DE) transcripts following a four hour freezing exposure at −6 °C relative to unfrozen control for each population. Up arrows indicate significantly upregulated transcripts, down arrows indicate significantly downregulated transcripts, and a tilde indicates contraregulated transcripts which are upregulated in one group, but downregulated in another. Areas indicate rank order of number of DE transcripts. A) Number of DE transcripts four hours following freezing between the White Sea and British Columbia populations. B) Number of DE transcripts four hours following freezing between the White Sea and British Columbia populations. C) Number of DE transcripts in the White Sea population (there were no DE transcripts before thawing). D) Number of DE transcripts in the British Columbia population.

### Identities of differentially expressed transcripts

The identities of the top differentially expressed transcripts following freezing were functionally very different between the two populations. The few transcripts that both populations upregulated immediately following freezing were putative transcription factors (annotated by Blast2Go as “proto-oncogenes”; Table 1). In the British Columbia population, the most noticeable pattern is the massive upregulation of transcripts during the freezing event (>9000 transcripts, Figures 3 and 4). This included several ribosomal RNAs, mitochondrially-encoded electron transport chain subunits, and histone methyltransferases and deacetylases (Supplementary Data 1). Among the downregulated transcripts includes several that Blast2Go annotated as “macrophage mannose receptors” which are functionally assigned the “carbohydrate binding” GO term (Supplementary Data 1). At four hours following freezing, this pattern has largely disappeared, with only a total of 46 significantly upregulated transcripts (Figure 3 and 4). These include several transcripts that Blast2Go identified as serine proteases, while the downregulated transcripts include one identified as “Peter Pan”, a transcript annotated as “macrophage mannose receptor”, and an INO80 complex subunit (Supplementary Data 1). By 28 hours following freezing, differential expression has increased again (Figure 3 and 4), with a total of 50 upregulated transcripts that BLAST to likely proteases and 13 that BLAST to ubiquitin activity, suggesting significant protein scavenging. By contrast, in the White Sea population, a large number of transcripts associated with heat shock proteins were the most highly upregulated transcripts following freezing (Table 2). Two transcripts that were identified as E3 ubiquitin ligases were also upregulated at both 4 and 28 hours following freezing (Supplementary Data 1). Four and 28 hours following freezing, the down-regulated transcripts included a variety of transcripts that perhaps most notably included several mitochondrially-encoded electron transport chain subunits (Supplementary Data 1) including NADH dehydrogenase subunits 1, 2, 4, 5, and 6, cytochrome b, ATP synthase F_0_ subunit 6, as well as cytochrome c oxidase subunits 1, 2, and 3 (Supplementary Data 1). No nuclear-encoded electron transport chain subunits were downregulated in our dataset.

**Table 1.**
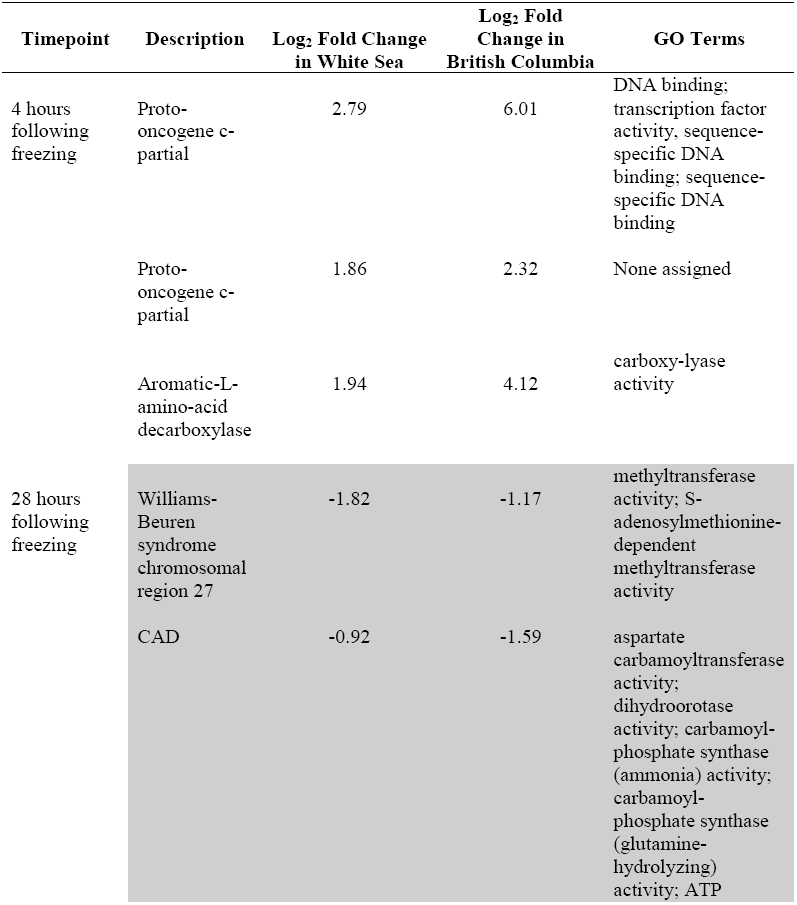
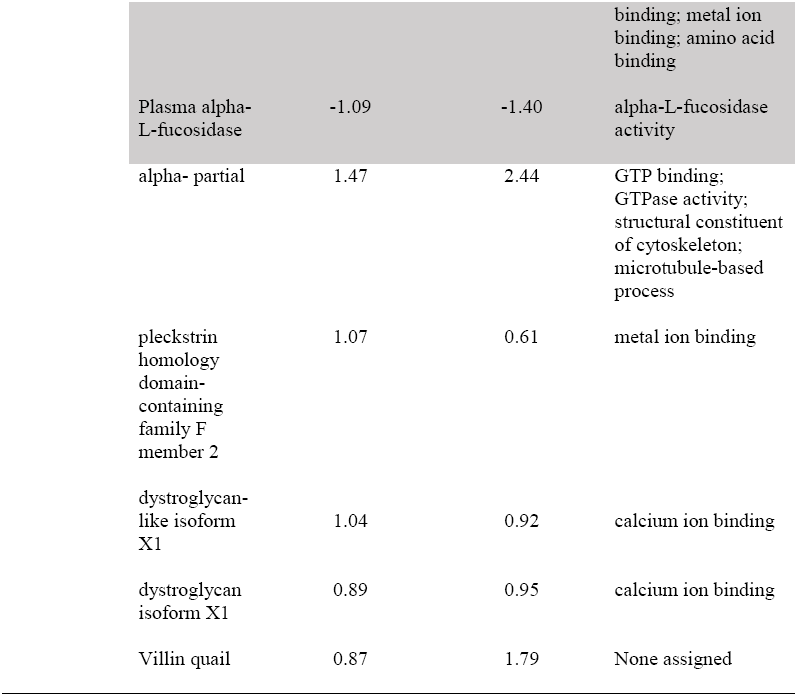
Transcripts that are significantly differentially regulated in both British Columbia and White Sea populations. GO terms are listed only for molecular function. Only the top 5 up and down-regulated transcripts are given (supplementary data 1 for all).

| Timepoint | Description | Log<sub>2</sub> Fold Change in White Sea | Log<sub>2</sub> Fold Change in British Columbia | GO Terms |
| --- | --- | --- | --- | --- |
| 4 hours following freezing | Proto-oncogene c-partial | 2.79 | 6.01 | DNA binding; transcription factor activity, sequence-specific DNA binding; sequence-specific DNA binding |
|  | Proto-oncogene c-partial | 1.86 | 2.32 | None assigned |
|  | Aromatic-L-amino-acid decarboxylase | 1.94 | 4.12 | carboxy-lyase activity |
| 28 hours following freezing | Williams-Beuren syndrome chromosomal region 27 | -1.82 | -1.17 | methyltransferase activity; S-adenosylmethionine-dependent methyltransferase activity |
|  | CAD | -0.92 | -1.59 | aspartate carbamoyltransferase activity; dihydroorotase activity; carbamoyl-phosphate synthase (ammonia) activity; carbamoyl-phosphate synthase (glutamine-hydrolyzing) activity; ATP |

|  |  |  |  |
| --- | --- | --- | --- |
|  |  |  | binding; metal ion binding; amino acid binding |
| Plasma alpha-L-fucosidase | -1.09 | -1.40 | alpha-L-fucosidase activity |
| alpha- partial | 1.47 | 2.44 | GTP binding; GTPase activity; structural constituent of cytoskeleton; microtubule-based process |
| pleckstrin homology domain-containing family F member 2 | 1.07 | 0.61 | metal ion binding |
| dystroglycan-like isoform X1 | 1.04 | 0.92 | calcium ion binding |
| dystroglycan isoform X1 | 0.89 | 0.95 | calcium ion binding |
| Villin quail | 0.87 | 1.79 | None assigned |

### Functional enrichment of molecular functions of differentially expressed transcripts

The functions of the transcripts similarly differed significantly between the two populations (Tables 2, 3, Figure 5). In the British Columbia population, we found significant enrichment of 134 GO terms after REVIGO filtering in the downregulated and 80 in the upregulated transcripts during freezing, including several electron transport chain-related terms in both sets (Figure 5). By four hours following freezing there was no significant enrichment of GO terms, and by 28 hours following freezing there was significant enrichment of 56 GO terms in the downregulated transcripts with only 15 in the upregulated transcripts (Figure 5). The most notable of these enriched terms included “UDP-glucose-6-dehydrogenase activity” and “scavenger receptor activity”. By contrast, in the White Sea population, as we had no significant differential expression of transcripts during freezing, we did not find any significantly enriched GO terms at this timepoint. While we did not find any significant enrichment of GO terms among upregulated transcripts in the White Sea population four hours following recovery, we did find a series of terms associated with the electron transport chain including “oxidoreductase activity”, “heme-copper terminal oxidase activity”, and “hydrogen ion transmembrane activity” that were significantly enriched among transcripts that were significantly downregulated at this timepoint (Figure 5). Interestingly, all of these downregulated terms were associated with transcripts that encoded only mitochondrially-encoded electron chain subunits. By 28 hours of recovery, the pattern had changed significantly with no enriched GO terms among the downregulated transcripts, but among the significantly upregulated transcripts the terms “glycerol transport”, “cellular water homeostasis”, and “water transport” were significantly enriched (which all were annotated as aquaporins).

**Figure 5.**
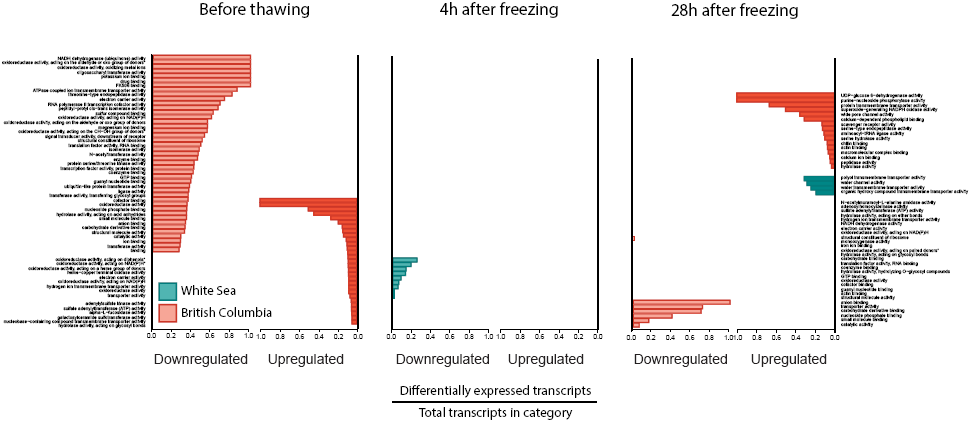
Significantly enriched GO terms at each timepoint in each population following a four hour freezing exposure at −6 °C relative to unfrozen control for each population as determined by Fisher’s exact test and REVIGO. Blue indicates terms enriched in the White Sea population, while red indicates terms enriched in the British Columbia population. Asterisks indicate terms with long names that have been shortened for space. All full terms are available in Supplementary Data 2.

**Table 2.** Transcripts that are significantly differentially regulated at both 4 and 28 hours following freezing in the White Sea population. GO terms are listed only for molecular function. Only the top 5 up and down-regulated transcripts are given (supplementary data 1 for all).

| Description | Log<sub>2</sub> Fold Change at 4 hours | Log<sub>2</sub> Fold Change at 28 hours | GO Names |
| --- | --- | --- | --- |
| heat shock partial | 2.79 | 6.01 | ATP binding |
| heat shock 68-like | 1.86 | 2.32 | ATP binding |
| heat shock 70 A1-partial | 1.94 | 4.12 | ATP binding |
| heat shock 70 partial | 1.82 | 1.17 | ATP binding |
| heat shock partial | 0.92 | 1.59 | ATP binding |
| gibberellin 20 oxidase | -1.09 | -1.40 | oxidoreductase activity |
| sodium-dependent nutrient amino acid transporter | -1.47 | -2.44 | symporter activity |
| BAI1-associated 3 | -1.07 | -0.61 | None assigned |
| Williams-Beuren syndrome chromosomal region 27 | -1.04 | -0.92 | methyltransferase activity; S-adenosylmethionine-dependent methyltransferase activity |
| None assigned | -0.89 | -0.95 | None assigned |

**Table 3.** Transcripts that are significantly differentially regulated at both 4 and 28 hours following freezing in the British Columbia population. GO terms are listed only for molecular function. Only the top 5 up and down-regulated transcripts are given (supplementary data 1 for all).

| Description | Log<sub>2</sub> Fold Change at 4 hours | Log<sub>2</sub> Fold Change at 28 hours | GO Names |
| --- | --- | --- | --- |
| Cytochrome partial | 6.61 | 5.62 | iron ion binding |
| Proto-oncogene c- partial | 6.31 | 2.52 | transcription factor activity, sequence-specific DNA binding |
| Proto-oncogene c- partial | 6.01 | 2.28 | transcription factor activity, sequence-specific DNA binding |
| techylectin-5A-like isoform X1 | 3.71 | 6.85 | carbohydrate binding |
| Proto-oncogene c- partial | 2.32 | 1.77 | None assigned |
| None assigned | -1.02 | -1.01 | None assigned |
| INO80 complex subunit | -0.74 | -0.59 | (Process: DNA repair) |
| Peter pan | -0.70 | -0.49 | rRNA binding |

Using pathview and Gage to examine significantly enriched gene sets based on KEGG orthology we found additional functional enrichment. In the British Columbia population, before thawing we found significant enrichment of the “spliceosome” term and underrepresentation of the “epithelial cell signaling in *Helicobacter pylori*” term. Four hours after thawing, we observed significant overrepresentation of the terms “ribosome biogenesis in eukaryotes” and “spliceosome”, and underrepresentation of “Parkinson’s disease”, “Alzheimer’s disease” and “oxidative phosphorylation”, although we note upon examination that the first two terms all indicated the same underrepresentation of electron transport chain subunits. There was no functional enrichment 28 hours following freezing in the British Columbia population. Finally, in the White Sea population we observed significant overrepresentation of terms including “ribosome biogenesis in eukaryotes”, “spliceosome”, and “bacterial invasion of epithelial cells” at 28 hours following freezing, but no terms were overrepresented at any other timepoints.

### Population differentiation

The estimated F_ST_ value (0.45; 9,164 loci passing filter) suggests little gene flow occurs between the two populations. Across all differentially expressed transcripts Tajima D estimates averaged −1.2 and −1.4 for White Sea and British Columbia respectively, whereas non-differentially expressed transcripts averaged −0.6 (T-test p-value <0.001) and −0.8 (T-test p-value <0.001) for White Sea and British Columbia respectively, suggesting selective sweeps on these differentially-expressed transcripts in each population (Table 4). Tajima’s D estimates for aquaporins in the White Sea population averaged −2.6 compared to the average across differently expressed transcripts of −1.42, indicative of a potential selective sweep in these transcripts (Table 4).

**Table 4.**
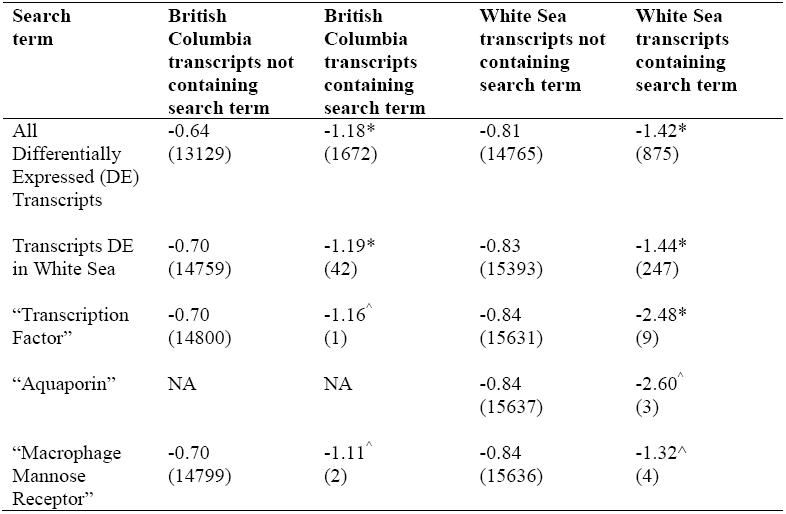
Tajima D estimates among transcripts. Estimates are averaged across transcripts that either contain or do not contain the search terms. No transcripts annotated as “aquaporins” had sufficient coverage to estimate Tajima D in the British Columbia population. The total number of transcripts passing filter for each test is in brackets. Numbers marked with an asterisk indicate a statistically-significant (t-test, P < 0.05) difference between the Tajima D estimates of transcripts containing the search term and those not. Numbers marked with a “^” indicates terms that had insufficient numbers of transcripts to run a t-test.

| **Search term** | **British Columbia transcripts not containing search term** | **British Columbia transcripts containing search term** | **White Sea transcripts not containing search term** | **White Sea transcripts containing search term** |
| --- | --- | --- | --- | --- |
| All Differentially Expressed (DE) Transcripts | -0.64<br>(13129) | -1.18*<br>(1672) | -0.81<br>(14765) | -1.42*<br>(875) |
| Transcripts DE in White Sea | -0.70<br>(14759) | -1.19*<br>(42) | -0.83<br>(15393) | -1.44*<br>(247) |
| “Transcription Factor” | -0.70<br>(14800) | -1.16^<br>(1) | -0.84<br>(15631) | -2.48*<br>(9) |
| “Aquaporin” | NA | NA | -0.84<br>(15637) | -2.60^<br>(3) |
| “Macrophage Mannose Receptor” | -0.70<br>(14799) | -1.11^<br>(2) | -0.84<br>(15636) | -1.32^<br>(4) |

## Discussion

Intertidal organisms constantly risk freezing through the winter. We find, consistent with our hypothesis, that the northern White Sea population of *S. balanoides* is significantly more freeze tolerant than the British Columbia population. To our knowledge, there is currently only one genome-sequenced freeze tolerant animal (the Antarctic midge *Belgica antarctica*; Kelley et al., 2014) and no genome-sequenced barnacles, so we assembled a *de novo* transcriptome to study transcriptional responses to freezing stress. In addition, the differing timelines of transcription suggest two very different sets of processes taking place in each population following a freezing event, which adds to our limited understanding of the molecular mechanisms of freeze tolerance. Finally, the low Tajima’s D values on these differentially expressed transcripts suggests that local adaptation to freezing has taken place in the White Sea population. Here we have shown the first study of transcriptional responses following a freezing stress in a non-insect, and demonstrated evidence of selection on transcripts associated with freeze tolerance in a natural population.

One of the most puzzling results we found is the massive upregulation of transcripts during the freezing exposure in the British Columbia population (Figures 3 and 4). There are three possibilities to explain this pattern: this represents genuine upregulation of transcription, degradation of mRNA causing greater numbers of smaller transcripts, or damage to translational machinery causing mRNA to accumulate. The first hypothesis of increased rates of *de novo* mRNA transcription is possible since the cytoplasm of cells generally remains unfrozen during a freezing exposure. However this is difficult to reconcile with the generally-accepted rate of transcription in invertebrates of about 2kb/minute (Ardehali & Lis, 2009), particularly at a subzero temperature where biochemical events would be expected to be proceed significantly more slowly. This upregulation is also strikingly absent in the White Sea population, suggesting increased transcription rates during freezing is not a general nor adaptive response to freezing stress. The second hypothesis of degradation of mRNA due to osmotic shock or direct freeze-induced damage is certainly possible, which could increase the total number of detected mRNA transcripts due to cleavage of larger molecules into several smaller ones. However, we note that the average transcript length of these upregulated transcripts is significantly larger (1761 bp vs. 1398 bp for the whole transcriptome), which is inconsistent with the possibility of mRNA degradation. We also note that, at least in *Arabidopsis thaliana*, low temperature tends to increase mRNA half-life (Chiba et al., 2013). The third hypothesis, cold-induced damage to ribosomal structure or translation initiation machinery, is possible since translation is associated with mRNA degradation (Bicknell & Ricci, 2017). At least in bacteria, the ribosome is particularly sensitive to cold shock and several cellular mechanisms for surviving cold shock are focused on ribosomal proteins (Phadtare, 2004). In addition, we note several ribosomal mRNAs are among the top-upregulated transcripts in this experimental group, as well as several transcripts associated with eukaryotic translation initiation machinery (Supplementary Data 1). Finally, we also note that at 4 hours following freezing, we find that the KEGG term “ribosome biogenesis” is significantly enriched among upregulated transcripts in the British Columbia population, suggesting repair or rebuilding of ribosomal subunits (Table S1). If the ribosomes are damaged and unable to process mRNA, we might expect a buildup of mRNA (Bicknell & Ricci, 2017). If this is true, we would expect that the White Sea population produces ribosomes that are more cold shock resistant. Further functional studies, in addition to further transcriptomic work in freeze tolerant animals, is clearly necessary to differentiate among these potential explanations.

The role(s) of heat shock proteins in freeze tolerance is not well understood, but upregulation of Hsp70 has been documented in response to repeated freezing in *B. antarctica* (Teets et al., 2011). While cold-induced denaturation of proteins has been well studied (Graziano, 2010; Todgham, Hoaglund, & Hofmann, 2007), during survivable freezing significant desiccation of cells is believed to occur (Storey, 1997; Teets & Denlinger, 2014). It is possible that this desiccation leads directly to protein denaturation through the disruption of the hydration shell around proteins during desiccating periods. In the more freeze tolerant White Sea population, we observed significant upregulation of several shock proteins following freezing—it is unclear whether these are involved with assisting in refolding of damaged proteins or are preparatory for a second stress event. By contrast, the less freeze tolerant population upregulated proteolysis-related transcripts (with significant enrichment of GO terms such as “serine endopeptidase activity” and “scavenger receptor”) following freezing, suggesting significant damage to protein structure has taken place and the only possible solution is to degrade these proteins. Our study points to the importance of maintenance of protein stability at low temperature in an ecological setting.

In addition to direct effects on protein structure, we also saw very different patterns of differential regulation of aquaporins between the two populations following freezing. In the White Sea population, we saw significant enrichment of GO categories associated with aquaporins, which we did not observe in the British Columbia population. This suggests that only the White Sea population is generally upregulating aquaporin transcription following freezing. It is believed that aquaporins aid in water transport out of the cell and that aquaglyceroporins may assist in transport of glycerol into the cell during freezing events (Philip et al., 2008; Storey & Storey, 2013; Yi et al., 2011). While significant glycerol accumulation is not associated with freeze tolerance in marine invertebrates (Storey & Storey, 2013), it is likely that managing water content in the cell is essential for freezing survival. Thus far aquaporins have been associated with freeze tolerance in the wood frog *Rana sylvatica* (Philip et al., 2008) and the goldenrod gall fly *Eurosta solidaginis* (Yi et al., 2011), and with this study we demonstrate their importance in a new class of freeze tolerant organisms, suggesting that this may be a generalized mechanism for freeze tolerance. In addition, we found very low Tajima’s D values in aquaporins that were upregulated following freezing (Table 4), suggesting that selective sweeps had taken place on the sequence of these putative aquaporins. Follow-up with functional studies, particularly employing the frog oocyte expression system, would provide interesting evidence and further structural information in this poorly understood system (Lind et al., 2017).

The role of antifreeze proteins in freeze tolerance is unclear and little is generally known about invertebrate antifreeze proteins (Davies, 2014; H. J. Kim et al., 2017). We found many transcripts (>200) that were annotated with the term “macrophage mannose receptors” that were upregulated following freezing in both populations. While we cannot definitively assign a function to these based solely on a transcriptomic study, we do note that these are C-type lectins, which are believed to be paralogous to Type II antifreeze proteins in fish (Graham, Lougheed, Ewart, & Davies, 2008; Ng & Hew, 1992). Indeed, when we examined the BLAST results, many of these sequences had high similarity to Type II antifreeze or ice structuring proteins in fish. Several freeze tolerant species are thought to produce antifreeze proteins to inhibit recrystallization of ice at low temperatures (Davies, 2014; Storey & Storey, 2013), although functional studies are necessary for confirming ice recrystallization inhibition activity. Antifreeze proteins are found broadly scattered across the domains of life, and appear to readily evolve (Davies, 2014). In addition, they appear to frequently have been transferred across species through horizontal gene transfer (Davies, 2014; Kiko, 2010). While this study was not designed to differentiate between the hypotheses of similar structure due to convergent evolution or by horizontal gene transfer, further sequencing studies may be able to address this point.

One of the most consistent results we found in both populations is downregulation of transcripts associated with the electron transport chain. There are two potential explanations for this downregulation—either transcription of mitochondrially-encoded transcripts was inhibited, or the number of mitochondria themselves were downregulated following a freezing stress. This is likely an adaptive response since a degraded or malfunctioning electron transport chain can cause significant oxidative stress, as can a sudden reduction in ATP demand as we might expect at low temperatures (Somero, Lockwood, & Tomanek, 2017). Reducing mitochondrial numbers as a way to reduce metabolic rate is a seasonal adaptation in the overwintering larvae of the moth *Gynaephora groenlandica* (Kukal, Duman, & Serianni, 1989), and this may be a similar phenomena. While we cannot distinguish between these explanations in this study, we hypothesize that reducing mitochondrial activity facilitates freeze tolerance in our barnacles and suggest studies isolating mitochondria to measure mitochondrial respiration and histology to determine mitochondrial numbers following freezing stress.

### Conclusions

Here we have presented the first transcriptional time series following freezing in a freeze tolerant barnacle. By comparing two populations that differ strongly in their freeze tolerance, we can control for genetic background and thereby infer the transcripts that are associated with greater freeze tolerance. We find transcriptional evidence for protein damage during a freezing event, and suggest that heat shock protein upregulation appears to be important for recovery from freezing. We also document aquaporin expression associated with freezing in a third freeze tolerant species, and describe a series of new putative antifreeze proteins. These transcripts were associated with selective sweeps in the White Sea population of barnacles, suggesting local adaptation to freezing conditions has occurred. Taken together, we shed new light on the mechanisms of adaptation to the poleward range margin in animals.

## Acknowledgements

The authors would like to acknowledge the Company of Biologists for travel funding to KEM, a University of Oklahoma startup grant and Nigel Fish for computing resources for KEM, as well as Taiwan-Russian collaboration funds from MOST, Taiwan (106-2923-B-001 −002 −MY3) and RFBR (grants17-54-52006 MNT_a, 18-04-00624, 15-29-02447 ofi_m) and Academia Sinica Senior Investigator Award (AS-IA-105-L03) to BKKC for funding for sequencing. We would like to thank the Hakai Institute and Mary O’Connor for collecting barnacles in British Columbia, as well as for access to temperature data. We would also like to acknowledge Yao-Fong Tsao and Vanessa Pei-Chen Tsai for laboratory assistance in sample preparation, and Dr. Mei-Yeh Lu for her invaluable advice for library preparation. Finally, we would like to thank Jantina Toxopeus and two anonymous referees for comments that have greatly improved the manuscript.

## Data accessibility

The original sequencing reads have been archived at the NCBI Short Read Archive under Bioproject number PRJNA496531 and will be made accessible on publication. The assembled and annotated *Semibalanus balanoides* transcriptome has deposited as a Transcriptome Shotgun Assembly at the NCBI database under the same number. Other data (read counts) have been deposited on the Open Science Framework at XXXXXX and will be made accessible upon publication.

## Author Contributions

KEM, AP, GK, and BKKC designed the research

BKKC, AP and GK collected the barnacles from the White Sea

KEM, AP, GK, and BKKC performed the laboratory research

KEM and ED analyzed the data

KEM wrote the first draft of the paper, AP, GK, BKKC, and ED contributed to writing

